# The Curli Accessory Protein CsgF Influences the Aggregation of Human Islet Amyloid Polypeptide

**DOI:** 10.1101/772392

**Authors:** Osmar Meza-Barajas, Isamar Aranda, Ashwag Binmahfooz, Alliosn Newell, Sajith Jayasinghe

**Affiliations:** Department of Chemistry and Biochemistry, California State University, San Marcos, CA 92096

## Abstract

Gram-negative bacteria, such as E. coli and Salmonella, contain proteinaceous, hair-like, cell surface filaments known as curli. Curli serve to facilitate cell-cell interactions and are essential for host cell colonization. Curli assembly involves six proteins, CsgA, CsgB, CsgC, CsgE, CsgF, and CsgG. CsgE and CsgF are thought to act as chaperones to help prevent the premature aggregation of CsgA and/or CsgB, and to help transport these proteins, through the outer-membrane protein CsgG, to the cell surface where they assemble to form Curli. It has been observed that CsgF is able to inhibit the aggregation of CsgA, the major protein component of Curli. This article describes CsgF’s ability to influence the aggregation of human islet amyloid polypeptide (hIAPP), an amyloidogenic polypeptide that is unrelated to Curli. In the presence of CsgF no increase in Thioflavin T fluorescence was observed for freshly solubilized hIAPP monitored as a function of time, suggesting that CsgF prevents the aggregation of hIAPP during the time period of observation. An analog of CsgF lacking the N-terminal unstructured region retained the ability to inhibit the aggregation of hIAPP. The nature of the CsgF-hIAPP interaction was probed via fluorescence quenching using a series of single cysteine mutants of CsgF labeled via the individual cysteine side chains with the fluorophore IAEDANS. In the presence of hIAPP, but not in the presence of the non-amyloidogenic rat islet amyloid polypeptide, the fluorophore attached to position of 23 of CsgF was found to be less exposed the quencher acrylamide suggesting that the interaction of hIAPP changes the solvent exposure of the N-terminus of CsgF. Taken together these data suggest that the structured region of CsgF, between residues 66 and 128, is involved in the protein’s interaction with hIAPP and that upon interaction structural changes make the N-terminus less solvent exposed.

## Background and Significance

Curli, proteinaceous filaments found on the outer surface of bacteria such as E. Coli and Salmonella, are involved in biofilm formation and bacterial attachment to surfaces (Austin, Sanders, Kay, & Collinson, 1998; Chapman et al., 2002; Collinson, Clouthier, Doran, Banser, & Kay, 1996; Römling, Bian, Hammar, Sierralta, & Normark, 1998; Sukupolvi et al., 1997). Investigation of Curli has shown that these fimbriae possess characteristics similar to amyloid fibers found in a number of diseases such as Alzheimer’s disease, Parkinson’s disease, and type II diabetes. Similar to amyloid, Curli were found to change the spectral properties of congo red and thioflavin T (ThT), as well as contain significant **β**-sheet character (Chapman et al., 2002). In contrast to amyloid found in disease, which are the result of protein misfolding, and are toxic to cells, Curli have a functional role in bacteria. Given these observations Curli have been classified as a functional amyloid: a protein aggregate with amyloid like properties that also serves a functional role. Curli assembly involves six proteins, Csg A, B, C, E, F, and G (Chapman et al., 2002). CsgA is the major protein component of Curli filaments while CsgB is thought to function as a nucleator of CsgA aggregation as well as to anchor CsgA to the outer surface of the bacterium (Bian & Normark, 1997; Hammar, Bian, & Normark, 1996). CsgC, CsgE, CsgF, and CsgG provide a variety of important support functions: CsgG, an outer-membrane channel, secretes the extracellular proteins to the outside milieu (Cao et al., 2014; Goyal et al., 2014); CsgC and CsgE, periplasmic proteins, prevent the intracellular aggregation of CsgA and/or CsgB (Evans et al., 2015; Gibson, White, Rajotte, & Kay, 2007; Taylor et al., 2011); while CsgF, found on the outer surface of bacteria, facilitates the extracellular assembly of CsgA into Curli. In the absence of CsgF, CsgA and CsgB do not localize to the cell surface and are secreted away (Chapman et al., 2002; Nenninger, Robinson, & Hultgren, 2009; Robinson, Ashman, Hultgren, & Chapman, 2006).

Three of the Curli accessory proteins, CsgC, CsgE, and CsgF, are able to influence the aggregation of CsgA in vitro. The in vitro aggregation of CsgA, as monitored using Thioflavin T (ThT) fluorescence, is inhibited by CsgC even when the two proteins are mixed at a very low molar ratio (1:500 CsgC:CsgA) suggesting that a single CsgC protein may be capable of interacting with more than one CsgA protein (Evans et al., 2015). CsgE was also found to prevent the aggregation of CsgA, when the two proteins were incubated together in an equimolar solution (Nenninger et al., 2011). The solution NMR structure of CsgE indicates a distinct distribution of positively and negatively charged residues, and it has been suggested that electrostatic interactions may play an important role in CsgE’s interaction with CsgA and its ability to prevent CsgA aggregation. The in vitro aggregation of CsgA is also inhibited by CsgF (Schubeis et al., 2018), but it is unclear if this inhibitory role is biologically relevant since, in vivo, CsgF is thought to function extracellularly to facilitate the proper assembly of CsgA in to Curli. The observation that CsgC, CsgE, and CsgF can inhibit amyloid formation by CsgA, and especially the ability of CsgC to inhibit amyloid formation by **α**-synuclein (Evans et al., 2015), point to the possibility that CsgC, CsgE, and CsgF could be used as tools to probe the details of protein-protein interactions that drive amyloid formation by CsgA, and perhaps amyloid formation in general. In this study the ability of CsgF to influence the aggregation of human Islet Amyloid Polypeptide (hIAPP), and the regions of CsgF that may be involved in interacting with hIAPP, was characterized. The results indicate that CsgF is able to inhibit the aggregation of hIAPP within the timescale of investigation and that residues within the N-terminus of CsgF experience changes in exposure in the presence hIAPP as compared to its absence.

## Material and Methods

Materials. pET21 vectors containing the inserted sequences for CsgF fused to the plasmid-encoded C-terminal hexahistidine tag were obtained from Genscript (Piscataway, NJ). E. Coli BL21(DE) expression competent cells, Bacterial Protein Extraction Reagent (B-PER), a C-terminal AntiHis antibody, 5-((((2-Iodoacetyl)amino)ethyl)amino)Naphthalene-1-Sulfonic Acid (IAEDANS), and Tris-(carboxyethyl) phosphine hydrochloride (TCEP) were obtained from ThermoFischer Scientific (Grand Island, NY). Hexafluoroisopropanol (HFIP) and ThT were obtained from Sigma-Aldrich (Milwaukee, WI). Synthetic wild-type human (hIAPP) and rat IAPP (rIAPP) were obtained from Bachem (King of Prussia, PA) or from PolyPeptide Laboratories (Torrence, CA).

Expression and Purification of C-terminal hexa-histidine tagged Salmonella Typhimurium CsgF. E. Coli BL21 (DE3) cells, transformed with the appropriate pET21 vector containing the sequence for Salmonella Typhimurium CsgF, fused to the plasmid encoded C-terminal hexa-histidine tag, were grown, at 37 ^0^C, to an OD (595 nm) of between 0.5 - 1. Protein production was induced with the addition of 1 mM IPTG, and the cells harvested by centrifugation after 16 hours of incubation at 26 ^0^C. Cells were resuspended in binding buffer (20 mM phosphate, 20 mM imidazole, 500 mM NaCl) and B-PER, frozen using liquid nitrogen and lysed using manual grinding with a motor and pestle. Unbroken cells and cell debris were removed by centrifugation (5000 x g) for 20 min at 4 ^0^C and histidine tagged protein was recovered from the lysate using a HiTrap TALON Crude Co^2+^ column (GE Healthcare, Piscataway, NJ) as described by the manufacturer. Resin bound protein was eluted from the column using elution buffer (20 mM phosphate, 250 mM imidazole, 500 mM NaCl) and subsequently desalted and buffer exchanged (in to 20 mM phosphate buffer) using a HiPrep 26/10 Desalting column (GE Healthcare, Piscataway, NJ). The purity of affinity purified histidine tagged CsgF was investigated using SDS PAGE and the presence of the histidine tag was confirmed by western blot using an Anti-His (C-Term)-HRP antibody (Life Technologies, Carlsbad, CA).

Preparation of IAPP. Lyophilized human or rat IAPP peptides were dissolved in HFIP to obtain clear solutions. Peptide concentrations were calculated and aliquots of peptide in HFIP were pipetted into 1.5 mL Eppendorf tubes, mixed with 500 μL of deionized distilled water, immediately frozen in liquid nitrogen, and lyophilized overnight. Dry lyophilized peptide was dissolved in appropriate buffer to yield 12.5 μM solutions prior to use in CD or fluorescence spectroscopy.

CD Spectroscopy. CD spectra were obtained using a Jaco 810 spectropolarimeter (Jasco Inc., Easton, MD). Measurements were taken every 0.5 nm at a scan rate of 50 nm/min with an averaging time of 1 s. All spectra were collected between 190 and 260 nm using a 2 mm path length quartz cuvette. Spectra were corrected by subtracting an appropriate background and are presented with intensity units of molar ellipticity.

ThT Fluorescence Assay For Protein Aggregation. Aggregation of IAPP was monitored using the fluorescence intensity increase of ThT. To each aggregation reaction, a sufficient amount of ThT to yield 25 μM (from 5 mM stock solution in deionized distilled water) was added immediately after dissolving IAPP in buffer, and real-time emission intensities were measured at 482 nm with excitation at 450 nm. Measurements were performed at room temperature with excitation and emission slit widths of 1 and 10 nm, respectively. Fluorescence measurements were taken using a FluoroMax-3 Spectro-fluorometer (Horiba Jobin Yvon Inc, Edison, NJ).

Fluorescence Labeling of CsgF Single Cysteine Mutants. Single cysteine mutants of CsgF, expressed and purified as described above, was incubated with 100-molar excess of TCEP at room temperature in pH 8, 20 mM phosphate buffer for one-half hour. 20-molar excess IAEDANS in dimethylsulfoxide was added and the mixture incubated at room temperature for 2 hours (or overnight at 4 ^0^C). Excess TCEP and IAEDANS label were removed using the desalting column as described above.

Fluorescence Quenching. Steady-state fluorescence quenching of IAEDANS-labeled CsgF single cysteine mutants were carried out on a FluoroMax-3 Spectrofluorometer (Horiba Jobin Yvon Inc, Edison, NJ) at room temperature. 400 μL of 12.5 μM labeled protein in 20 mM phosphate buffer was titrated with 1 μL of quencher (Acrylamide) and IAEDANS fluorescence emission spectra were recorded from 400 - 600 nm using an excitation of 336 nm (with excitation and emission and slits widths of 5 nm). Accessibility of the quencher to the fluorophore was qualified using the Stern-Volmer quenching constant (K_sv_) obtained by fitting normalized fluorescence intensities to the Stern-Volmer equation F_0_/F = 1+ K_sv_[Q], where F_0_ and F are the fluorescence intensities in the absence and presence of quencher respectively, and [Q] is the quencher concentration. Intensities were determined at the wavelength of maximum emission in the absence of quencher.

## Results

### CsgF Inhibits The Aggregation of hIAPP

To determine the influence of CsgF on the aggregation of hIAPP, the real time fluorescence intensities of ThT at 482 nm was measured in the absence and presence of CsgF. In the absence of CsgF freshly prepared hIAPP exhibits a sigmoidal increase in ThT intensity (Figure 2A, open squares) with a time to half-maximal fluorescence intensity (t50) of approximately 9 (± 4) hrs. During the time in which aggregation was monitored an 11 (± 5) fold increase in ThT fluorescence intensity was observed. The CD spectrum of freshly dissolved hIAPP exhibited a peak with negative ellipticity at ∼ 198 nm indicative of a predominantly unstructured backbone, while the spectrum obtained after the increase in ThT intensity contained a peak with negative ellipticity at ∼ 218 nm, indicative of a predominantly **β**-sheet backbone structure (Figure 2C). These observations of freshly dissolved hIAPP are similar to those described in the literature for hIAPP aggregation (Kapurniotu, 2001; Padrick & Miranker, 2002). Incubating hIAPP with CsgF in a 1:1 mole ratio completely abolished the increase in ThT intensity (Figure 2A, closed squares), and no increase in ThT intensity was observed for the sample containing CsgF during the time period in which aggregation was monitored for hIAPP alone. Unlike in the case of hIAPP alone no significant change was observed between the CD spectrum obtained immediately after mixing CsgF with hIAPP and the spectrum obtained at the end of the ThT time course. The ratio of intensities at 208 and 222 nm are 1.28 for both spectra indicating the absence of any significant secondary structure change.

### In the Presence of hIAPP the Fluorophore at the N-terminus of CsgF is Protected from Quenching

To determine the nature of the CsgF-IAPP interaction, five single cysteine mutants of CsgF, labeled with the fluorophore IAEDANS, were exposed to a quencher in the absence and presence of hIAPP. Fluorescence intensities (F_0_/F) obtained as a function of increasing quencher concentrations were plotted to determine the Stern-Volmer quenching constant, K_sv_ (Figure 3A), which can be used to gauge the exposure of the quencher to the fluorophore. For the protein alone, CsgF labeled at positions 23, 44, 66, 97, and 123 exhibited K_sv_ values of 11.00 ± 0.38, 7.76 ± 0.18, 8.80 ± 0.50, 8.37 ± 0.40, and 12.33 ± 0.48 respectively. K_sv_ values obtained for the fluorophore placed close to the N- and C-terminal residues (23 and 123 respectively) are much higher than those observed for the fluorophore located on residues 44, 66, and 97, towards the middle of the amino acid sequence. These higher K_sv_ values suggested that residues in the N- and C-terminal regions are much more accessible to the quencher than those in the middle of the protein. This observation is in agreement with the solution NMR structure of E. Coli CsgF which exhibits unstructured N- and C-terminal regions with a well-defined **α**-helix and four stranded **β**-sheet (Figure 1B) occupying the central region of the protein (Schubeis et al., 2018). Although there does not appear to be a well-defined globular fold to CsgF, it is possible that residues in the **α**-helix and **β**-sheet regions of the protein are less solvent accessible than residues in the terminal regions, due to interactions between the structured regions.

**Figure 1:**
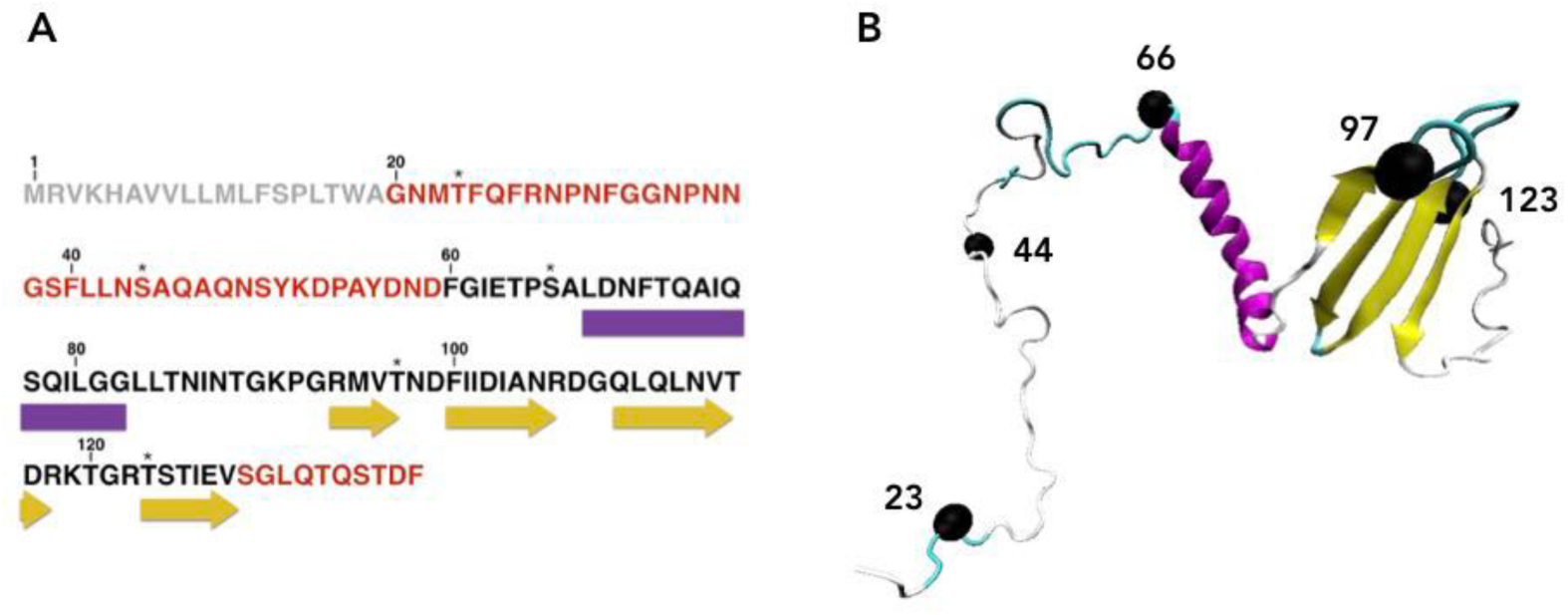
(A). Full length amino acid sequence of Salmonella typhimurium CsgF (P0A202). Regions of secondary structure identified in the NMR structure of Escherichia Coli CsgF in DHPC micelles (PDB ID: 5M1U) are represented using purple rectangles (**α**-helices) and yellow arrows (**β**-strands). The N- and C-terminal regions with no assigned secondary structure, and predicted to be unstructured (A. Green et al., 2016), are shown in red. Residues changed to cysteines to produce single cysteine mutants (CsgF T23C, CsgF S44C, CsgF S66C, CsgF T97C and CsgF T123C) used in this study are indicated by an asterisk. The N-terminal 19 residue signal sequence is shown in light grey. (B). NMR structure of E. Coli CsgF in DHPC micelles (PDB ID: 5M1U). Residues 23, 44, 66, 97, and 123 mutated to cysteines in this study correspond to residues 27, 49, 80 and 106 of the NMR structure and are rendered as black spheres. For consistency residues are numbered according to the full length sequence.

**Figure 2:**
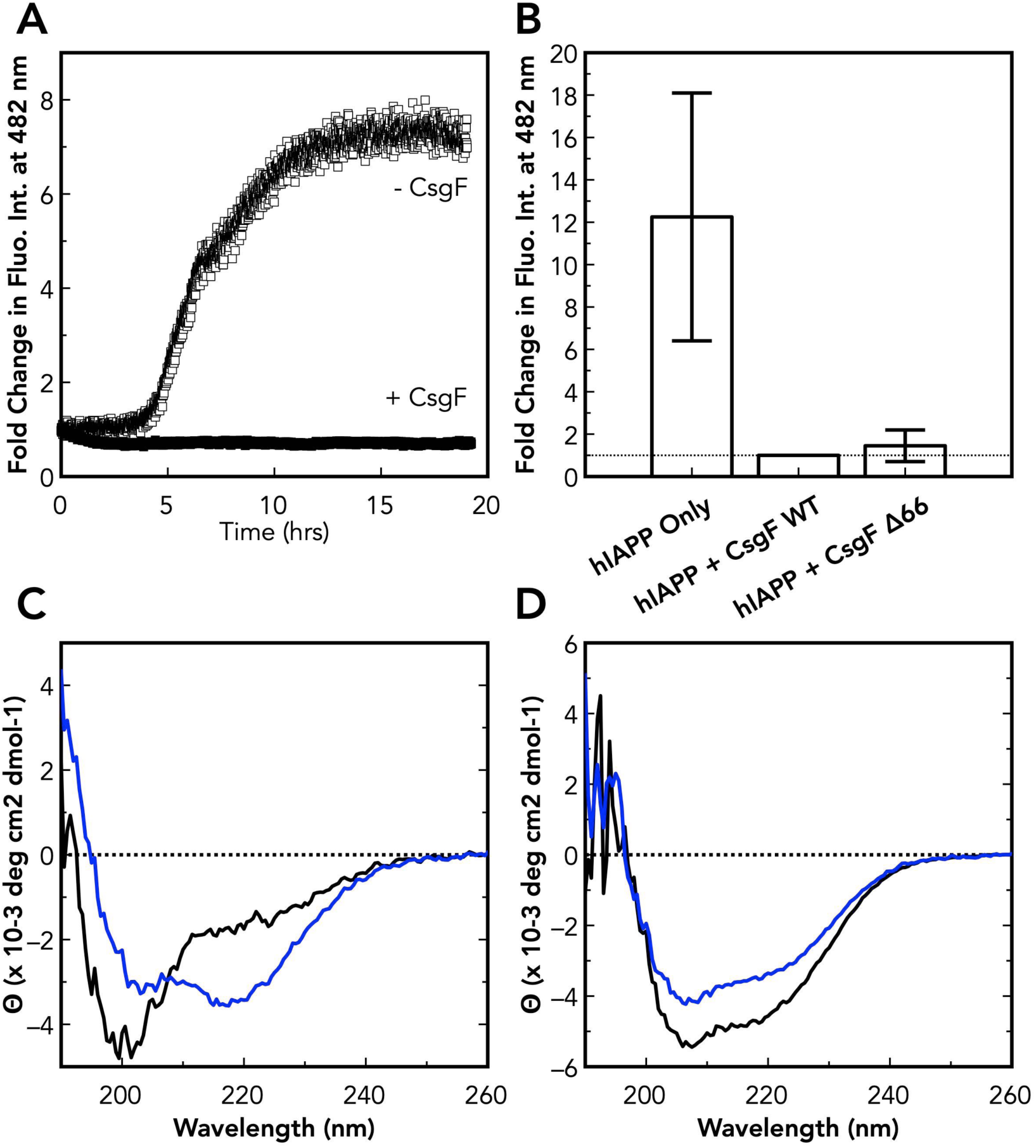
CsgF inhibits the aggregation of hIAPP. Aggregation of 12.5 μM hIAPP in 20 mM phosphate buffer was monitored using ThT fluorescence intensity at 482 nm. In order to facilitate comparison between different samples ThT intensity immediately after mixing CsgF with hIAPP is normalized to 1. (A) In the absence of CsgF (open squares) there is significant increase in ThT florescence intensity demonstrating hIAPP aggregation. In the presence of CsgF (closed squares) no increase in ThT fluorescence intensity is observed indicating an absence of hIAPP aggregation. (B) Average fold change in ThT fluorescence intensity in the absence, and presence of CsgF, and in the presence of CsgFΔ66. The average ThT fluorescence intensity increase in the absence of CsgF is significantly different than the intensity increase observed in the presence of CsgF. Error bars represent the standard deviation of three individual experiments. In order to be able to compare the differences in hIAPP aggregation in the absence and presence of CsgF one hIAPP sample was divided into two. Both hIAPP samples, one without CsgF, and the other with, were monitored simultaneously on two spectrofluorometers. (C) CD spectra of 12.5 μM hIAPP (in the absence of CsgF) immediately after rehydrating in buffer (black line) and at the end of the ThT time course (∼ 20 hrs, blue line). CD spectrum of hIAPP immediately after rehydrating in buffer are indicative of an unordered structure. The spectrum obtained after the ThT intensity had increased contains a negative intensity at ∼ 218 nm and is indicative of the appearance of **β**-sheet secondary structure as expected for the aggregation of hIAPP. (D) CD spectra obtained immediately after mixing 12.5 μM CsgF with 12.5 μM hIAPP (black line) and art the end of the ThT time course. Although the spectrum at the end of the time course exhibits less negative intensity between 200-240 nm there does not appear to be any significant change in secondary structure. The ratio of intensity at 208 to 222 nm for the spectrum obtained at the end of the time course remains the same (1.28) as the ratio obtained from the spectrum immediately after mixing CsgF with hIAPP.

**Figure 3:**
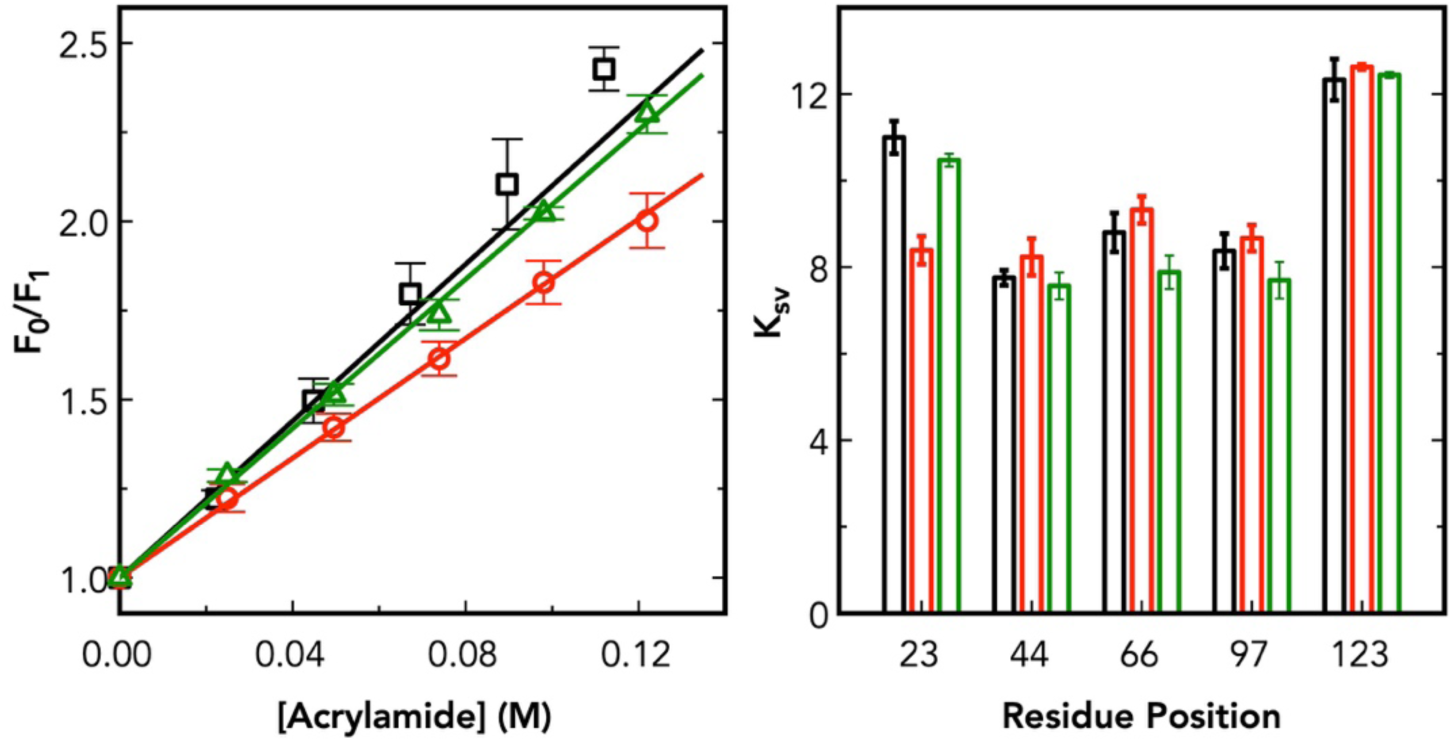
hIAPP protects fluorophore attached to position 23 of CsgF from quenching by acrylamide. (A) Stern-Volmer plot of acrylamide quenching of 12.5 μM IAEDANS labeled CsgF alone (black), and in the presence of 12.5 μM hIAPP (red) and 12. 5 μM rIAPP (green) in 20 mM phosphate buffer at room temperature. Stern-Volmer quenching constants were determined using linear fits to the data. (B) Stern-Volmer quenching constants for CsgF labeled at positions 23, 44, 66, 97, and 123 (black bars), and in the presence of hIAPP (red bars), and in the presence of rIAPP (green bars). Error bars represent the standard deviation of three individual experiments.

In the presence of equimolar hIAPP a significantly lower K_sv_ value, 8.39±0.32, was obtained for the fluorophore at position 23 suggesting a protection of the fluorophore from the quencher. Quenching experiments were also carried out in the presence of rat islet amyloid polypeptide (rIAPP) which differs from hIAPP by only six residues and does not aggregate to form fibrils. The K_sv_ of 10.47±0.15 observed for the fluorophore at position 23 in the presence of rIAPP is similar to that obtained for the protein alone and suggests that the reduction in K_sv_ observed in the presence of hIAPP is a result of a specific interaction between CsgF and hIAPP. No significant change in K_sv_ values were observed for fluorophores placed at positions 44, 66, 97, or 123 of CsgF in the presence of hIAPP or rIAPP (Figure 3B) indicating that the interaction with hIAPP only protects fluorophores placed close to the N-terminus of CsgF.

### The N-terminal residues of CsgF are not needed to inhibit the aggregation of hIAPP

To determine if the N-terminal region of CsgF plays a role in the protein’s ability to inhibit the aggregation of hIAPP, aggregation was monitored in the presence of an analog of CsgF lacking the first 66 residues (CsgFD66). Freshly rehydrated hIAPP incubated in the presence of CsgFD66 exhibits a slight (∼ 2-fold) increase in ThT fluorescence (Figure 2B). The increase in ThT fluorescence is significantly less than that observed for hIAPP alone and thus it appears that CsgFD66 is still able to appreciably inhibit the aggregation if hIAPP, and that the N-terminal unstructured region of CsgF is not crucial to this activity.

## Discussion

CsgF plays a crucial role in the formation of bacterial Curli with the protein required for the proper assembly of the major Curli subunit protein CsgA into fibrils on the outer surface of bacteria (Chapman et al., 2002; Nenninger et al., 2009; Robinson et al., 2006). Although the exact mechanism of the protein is not known, it has been speculated that CsgF could help to bind and serve as a template for folding and aggregation of CsgA and CsgB, and additionally perhaps also function to anchor CsgA/CsgB to the outer surface of the bacterium (Schubeis et al., 2018). Interestingly CsgF influences the aggregation of CsgA and CsgB in vitro. ThT aggregation assays have shown that the protein inhibits the aggregation of CsgA while stabilizing and aiding the aggregation of CsgB (Schubeis et al., 2018). ThT fluorescence data presented here suggests that CsgF is also able to inhibit the in vitro aggregation of hIAPP (Figure 2A). hIAPP is a 37-residue amyloidogenic protein implicated in type II diabetes which has been shown to form in-register parallel **β**-sheet structures upon aggregation (Bedrood et al., 2012). Although intriguing, the ability of a Curli accessory protein to inhibit the aggregation of an amyloidogenic protein unrelated to Curli formation is not unique. ThT aggregation studies have shown that CsgC is able to prevent the in vitro aggregation of human **α**-synuclein, an amyloidogenic protein implicated in Parkinson’s disease (Evans et al., 2015). The ability of CsgC to prevent protein aggregation appears to be specific since it was unable to prevent the aggregation of the Alzheimer’s A**β**_42_ peptide even at a 1:1 (CsgC:A**β**_42_) molar ratio. Based on a comparison of the primary structure of CsgA and **α**-synuclein, it has been suggested that these two proteins interact with CsgC via a common interaction motif (Evans et al., 2015). In contrast, there is no discernible region of sequence similarity between CsgA and hIAPP, suggesting that the ability of CsgF to inhibit the aggregation of hIAPP does not involve the recognition of a specific amino acid sequence.

Characterization of CsgF has shown that a significant portion of the protein is unstructured (A. Green et al., 2016; Schubeis et al., 2018). The solution NMR structure shows a lack of secondary structure for the N-terminal forty residues as well as the approximately ten residues at the C-terminus (Schubeis et al., 2018). Immediately following the N-terminal unstructured region is a 21 residue **α**-helix and a four stranded **β**-sheet (Figure 1). Fluorescence quenching experiments carried out to probe the region of CsgF that interacts with hIAPP indicates that upon mixing hIAPP with CsgF, at equimolar concentrations, only the fluorophore placed at position 23 of CsgF becomes protected from the quencher (Figure 3A). This suggests that interaction of hIAPP with CsgF either occurs close to the N-terminus of CsgF, or that the interaction causes a structural change in CsgF causing the N-terminus to become less solvent exposed. The latter hypothesis is more likely given the observation that a CsgF analog lacking the N-terminal 66 residues (CsgFΔ66) was still able to significantly inhibit the aggregation of hIAPP (Figure 1B). Analysis of the CD spectrum of CsgFΔ66 (Figure 4) using DichroWeb (Whitmore & Wallace, 2004) indicates a secondary structure distribution of approximately 15 % **α**-helix, 30% **β**-strand, 20% turn and 35% of the protein being unstructured. This distribution compares favorably with the secondary structures observed for the solution NMR structure for this region (approximately 21% a-helix, 38% b-strand, 11% turn, and 29% unstructured) suggesting that removal of the N-terminal unstructured region does not alter the overall secondary structure distribution of the remaining amino acid sequence. These observations support the hypothesis that the structured region of CsgF, approximately residues 66-128, are involved in the interaction with hIAPP and that upon interaction the unstructured N-terminus of CsgF becomes less solvent exposed.

**Figure 4:**
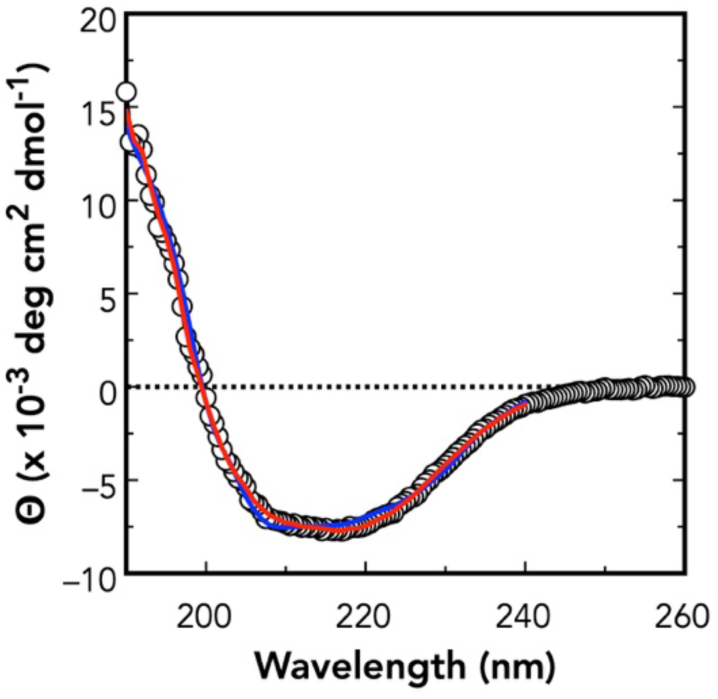
CD Spectrum of CsgFΔ66 in 20 mM phosphate buffer. The spectrum was deconvoluted using DICHROWEB and CsgFΔ66 was estimated to contain approximately 15 % **α**-helix, 30% **β**-strand, and 20% turn secondary structure distribution with 35% of the protein being unstructured. Fitted spectra obtains via DICHROWEB are shown in red (CDSSTR) and blue (CONTIN).

Although the fluorophore placed at position 23 of CsgF experienced a significant protection from the quencher in the presence of hIAPP, quenching experiments carried out in the presence of rIAPP do not show any protection from the quencher regardless of the placement the fluorophore in CsgF. rIAPP differs from hIAPP by only six residues with the majority of the difference contained between residues 23 and 29. Residues 20-29 of hIAPP are regarded as the most critical for amyloid formation (Goldsbury et al., 2000; Moriarty & Raleigh, 1999; Rhoades, Agarwal, & Gafni, 2000; Tenidis et al., 2000) and the inability of rIAPP to aggregate to form amyloid has been attributed to the presence of three proline residues within this region (J. Green, 2003). It is conceivable that these proline residues prevent the interaction of rIAPP with CsgF leading to the differences in quenching observed at position 23.

The observation that CsgF can influence the aggregation of hIAPP described here adds to the growing body of evidence indicating that the Curli accessory proteins CsgC, CsgE, and CsgF can modulate the aggregation of proteins to form amyloid. Future studies aimed at more clearly identifying the regions and residues of CsgF involved in interacting with hIAPP as well as CsgA could possibly help determine the common features of amyloid formation in general and the nature of CsgA aggregation to form Curli.

## Acknowledgements

This work was supported by National Institutes of Health Grant 1R15GM123430-01 (to S.J.).

## References

Austin, J. W., Sanders, G., Kay, W. W., & Collinson, S. K. (1998). Thin aggregative fimbriae enhance Salmonella enteritidis biofilm formation. FEMS Microbiology Letters, 162(2), 295–301.

Bedrood, S., Li, Y., Isas, J. M., Hegde, B. G., Baxa, U., Haworth, I. S., & Langen, R. (2012). Fibril structure of human islet amyloid polypeptide. Journal of Biological Chemistry, 287(8), 5235–5241. http://doi.org/10.1074/jbc.M111.327817

Bian, Z., & Normark, S. (1997). Nucleator function of CsgB for the assembly of adhesive surface organelles in Escherichia coli. The EMBO Journal, 16(19), 5827–5836. http://doi.org/10.1093/emboj/16.19.5827

Cao, B., Zhao, Y., Kou, Y., Ni, D., Zhang, X. C., & Huang, Y. (2014). Structure of the nonameric bacterial amyloid secretion channel. Proceedings of the National Academy of Sciences of the United States of America, 111(50), E5439–44. http://doi.org/10.1073/pnas.1411942111

Chapman, M. R., Robinson, L. S., Pinkner, J. S., Roth, R., Heuser, J., Hammar, M., et al. (2002). Role of Escherichia coli curli operons in directing amyloid fiber formation. Science, 295(5556), 851–855. http://doi.org/10.1126/science.1067484

Collinson, S. K., Clouthier, S. C., Doran, J. L., Banser, P. A., & Kay, W. W. (1996). Salmonella enteritidis agfBAC operon encoding thin, aggregative fimbriae. Journal of Bacteriology, 178(3), 662–667. http://doi.org/10.1128/jb.178.3.662-667.1996

Evans, M. L., Chorell, E., Taylor, J. D., Åden, J., Götheson, A., Li, F., et al. (2015). The bacterial curli system possesses a potent and selective inhibitor of amyloid formation. Molecular Cell, 57(3), 445–455. http://doi.org/10.1016/j.molcel.2014.12.025

Gibson, D. L., White, A. P., Rajotte, C. M., & Kay, W. W. (2007). AgfC and AgfE facilitate extracellular thin aggregative fimbriae synthesis in Salmonella enteritidis. Microbiology (Reading, England), 153(Pt 4), 1131–1140. http://doi.org/10.1099/mic.0.2006/000935-0

Goldsbury, C., Goldie, K., Pellaud, J., Seelig, J., Frey, P., Müller, S. A., et al. (2000). Amyloid fibril formation from full-length and fragments of amylin. Journal of Structural Biology, 130(2-3), 352–362. http://doi.org/10.1006/jsbi.2000.4268

Goyal, P., Krasteva, P. V., Van Gerven, N., Gubellini, F., Van den Broeck, I., Troupiotis-Tsaïlaki, A., et al. (2014). Structural and mechanistic insights into the bacterial amyloid secretion channel CsgG. Nature, 516(7530), 250–253. http://doi.org/10.1038/nature13768

Green, A., Pham, N., Osby, K., Aram, A., Claudius, R., Patray, S., & Jayasinghe, S. A. (2016). Are the curli proteins CsgE and CsgF intrinsically disordered? Intrinsically Disordered Proteins, 4(1), e1130675. http://doi.org/10.1080/21690707.2015.1130675

Green, J. (2003). Full-length Rat Amylin Forms Fibrils Following Substitution of Single Residues from Human Amylin. Journal of Molecular Biology, 326(4), 1147–1156. http://doi.org/10.1016/S0022-2836(02)01377-3

Hammar, M., Bian, Z., & Normark, S. (1996). Nucleator-dependent intercellular assembly of adhesive curli organelles in Escherichia coli. Proceedings of the National Academy of Sciences of the United States of America, 93(13), 6562–6566.

Kapurniotu, A. (2001). Amyloidogenicity and cytotoxicity of islet amyloid polypeptide. Biopolymers, 60(6), 438–459. http://doi.org/10.1002/1097-0282(2001)60:6<438::AID-BIP10182>3.0.CO;2-A

Moriarty, D. F., & Raleigh, D. P. (1999). Effects of sequential proline substitutions on amyloid formation by human amylin20-29. Biochemistry, 38(6), 1811–1818. http://doi.org/10.1021/bi981658g

Nenninger, A. A., Robinson, L. S., & Hultgren, S. J. (2009). Localized and efficient curli nucleation requires the chaperone-like amyloid assembly protein CsgF. Proceedings of the National Academy of Sciences of the United States of America, 106(3), 900–905. http://doi.org/10.1073/pnas.0812143106

Nenninger, A. A., Robinson, L. S., Hammer, N. D., Epstein, E. A., Badtke, M. P., Hultgren, S. J., & Chapman, M. R. (2011). CsgE is a curli secretion specificity factor that prevents amyloid fibre aggregation. Molecular Microbiology, 81(2), 486–499. http://doi.org/10.1111/j.1365-2958.2011.07706.x

Padrick, S. B., & Miranker, A. D. (2002). Islet amyloid: phase partitioning and secondary nucleation are central to the mechanism of fibrillogenesis. Biochemistry, 41(14), 4694–4703. http://doi.org/10.1021/bi0160462

Rhoades, E., Agarwal, J., & Gafni, A. (2000). Aggregation of an amyloidogenic fragment of human islet amyloid polypeptide. Biochimica Et Biophysica Acta (BBA) - Protein Structure and Molecular Enzymology, 1476(2), 230–238. http://doi.org/10.1016/S0167-4838(99)00248-4

Robinson, L. S., Ashman, E. M., Hultgren, S. J., & Chapman, M. R. (2006). Secretion of curli fibre subunits is mediated by the outer membrane-localized CsgG protein. Molecular Microbiology, 59(3), 870–881. http://doi.org/10.1111/j.1365-2958.2005.04997.x

Römling, U., Bian, Z., Hammar, M., Sierralta, W. D., & Normark, S. (1998). Curli fibers are highly conserved between Salmonella typhimurium and Escherichia coli with respect to operon structure and regulation. Journal of Bacteriology, 180(3), 722–731.

Schubeis, T., Spehr, J., Viereck, J., Köpping, L., Nagaraj, M., Ahmed, M., & Ritter, C. (2018). Structural and functional characterization of the Curli adaptor protein CsgF. FEBS Letters, 592(6), 1020–1029. http://doi.org/10.1002/1873-3468.13002

Sukupolvi, S., Lorenz, R. G., Gordon, J. I., Bian, Z., Pfeifer, J. D., Normark, S. J., & Rhen, M. (1997). Expression of thin aggregative fimbriae promotes interaction of Salmonella typhimurium SR-11 with mouse small intestinal epithelial cells. Infection and Immunity, 65(12), 5320–5325.

Taylor, J. D., Zhou, Y., Salgado, P. S., Patwardhan, A., McGuffie, M., Pape, T., et al. (2011). Atomic resolution insights into curli fiber biogenesis. Structure (London, England: 1993), 19(9), 1307–1316. http://doi.org/10.1016/j.str.2011.05.015

Tenidis, K., Waldner, M., Bernhagen, J., Fischle, W., Bergmann, M., Weber, M., et al. (2000). Identification of a penta- and hexapeptide of islet amyloid polypeptide (IAPP) with amyloidogenic and cytotoxic properties. Journal of Molecular Biology, 295(4), 1055–1071. http://doi.org/10.1006/jmbi.1999.3422

Whitmore, L., & Wallace, B. A. (2004). DICHROWEB, an online server for protein secondary structure analyses from circular dichroism spectroscopic data. Nucleic Acids Research, 32(Web Server issue), W668–73. http://doi.org/10.1093/nar/gkh371

